# Single-Cell Peripheral Immunoprofiling of Lewy Body Disease in a Multi-site Cohort

**DOI:** 10.1101/2024.01.30.578082

**Authors:** Thanaphong Phongpreecha, Kavita Mathi, Brenna Cholerton, Eddie J. Fox, Natalia Sigal, Camilo Espinosa, Momsen Reincke, Philip Chung, Ling-Jen Hwang, Chandresh R. Gajera, Eloise Berson, Amalia Perna, Feng Xie, Chi-Hung Shu, Debapriya Hazra, Divya Channappa, Jeffrey E. Dunn, Lucas B. Kipp, Kathleen L. Poston, Kathleen S. Montine, Holden T. Maecker, Nima Aghaeepour, Thomas J. Montine

**Affiliations:** Department of Pathology, Stanford University, CA, USA; Department of Anesthesiology, Perioperative and Pain Medicine, Stanford University, CA, USA; Department of Biomedical Data Science, Stanford University, CA, USA; Institute for Immunity, Transplantation and Infection, Stanford University, CA, USA; Department of Pediatrics, Stanford University, CA, USA; Department of Neurology and Neurological Sciences, Stanford University, CA, USA

**Keywords:** Parkinson’s Disease, Alzheimer’s Disease, Biomarkers, Dementia, Inflammation

## Abstract

Studies implicated peripheral organs involvement in the development of Lewy body disease (LBD), a spectrum of neurodegenerative diagnoses that include Parkinson’s Disease (PD) without or with dementia (PDD) and dementia with Lewy bodies (DLB). This study characterized peripheral immune responses unique to LBD at single-cell resolution. Peripheral mononuclear cell (PBMC) samples were collected from sites across the U.S. The diagnosis groups comprise healthy controls (HC, n=164), LBD (n=132), Alzheimer’s disease dementia (ADD, n=98), other neurodegenerative disease controls (NDC, n=21), and immune disease controls (IDC, n=14). PBMCs were activated with three stimulants, stained by surface and intracellular signal markers, and analyzed by flow cytometry, generating 1,184 immune features. Our model classified LBD from HC with an AUROC of 0.90±0.06. The same model distinguished LBD from ADD, NDC, IDC, or other common conditions associated with LBD. Model predictions were driven by pPLCγ2, p38, and pSTAT5 signals from specific cell populations and activations.

## Introduction

Lewy Body Disease (LBD) comprises a spectrum of clinically and pathologically overlapping conditions: Dementia with Lewy Bodies (DLB) and Parkinson’s Disease (PD) with or without Dementia (PDD)^1–5^. Human genetic, biochemical, and pathological evidence, as well as experimental models, support involvement not only by neuroinflammation^6–8^ but also a peripheral immune response in the initiation and/or progression of LBD^6,9–13^. While there is intense interest in the systemic origins of pathologic alpha-synuclein, the role of the peripheral immune system in LBD remains unclear. One possibility is that subsets of peripheral immune cells migrate into the brain and consequently play a direct role in neurodegeneration^14^. Alternatively, peripheral immune cells may serve as biomarkers of an inherited or acquired trait shared by both peripheral and brain immune cells without peripheral cells directly contributing to neurodegeneration. Past research has explored peripheral blood mononuclear cells (PBMCs) as a platform to gain insights into the development of LBD with a focus on changes in the proportion of specific cell types^15–20^ or concentration of intercellular signals such as interleukins (ILs)^21–24^. While alteration of intracellular signaling in PBMCs of cognitively impaired or Alzheimer’s Disease (AD) patients has been explored previously^25–29^, only a handful of studies have profiled PBMC intracellular signaling for LBD^30–32^. Moreover, most of these investigations of PBMCs in LBD have been limited by small sample sizes, single cohorts, bulk analysis, and lack of disease controls to determine non-specific changes related to neurodegenerative diseases or immune-mediated diseases.

This study sought to address several of these limitations through a rigorous profiling of peripheral immune responses by PBMCs from 429 age- and sex-matched multisite research participants diagnosed with LBD, other neurodegenerative diseases (NDC), or healthy controls (HC). Fourteen additional samples also were obtained from patients at a single site who were diagnosed with autoimmune disease. Samples were unstimulated or activated with three different canonical immune stimulants to gain functional insight and then assayed with a panel of markers that resolved 37 different cell types and the intracellular signaling pathways that were selected to encompass those previously implicated by genetic risk and their associated pathways^33–36^.

## Results

### Overview of the Cohort and Immune Features

Samples were from individuals with one of these clinical diagnoses: healthy controls (HC), LBD, ADD, other neurodegenerative disease controls (NDC), or autoimmune disease controls (IDC). All diagnosis groups were exclusive, *e.g.* no patients were diagnosed with both LBD and AD. Each individual’s PBMCs were stimulated with LPS, IFNa, IL6, or unstimulated, followed by staining and measurement of cell type-specific abundance and intracellular signaling (see **Methods Section**), including Lamp2, p38, pPLCγ2, pS6, pSTAT1, pSTAT5, and Rab5. After cell type gating, there were 1,184 immune features total in each of the 429 individual PBMC samples (**Fig. 1A**).

**Figure 1.**
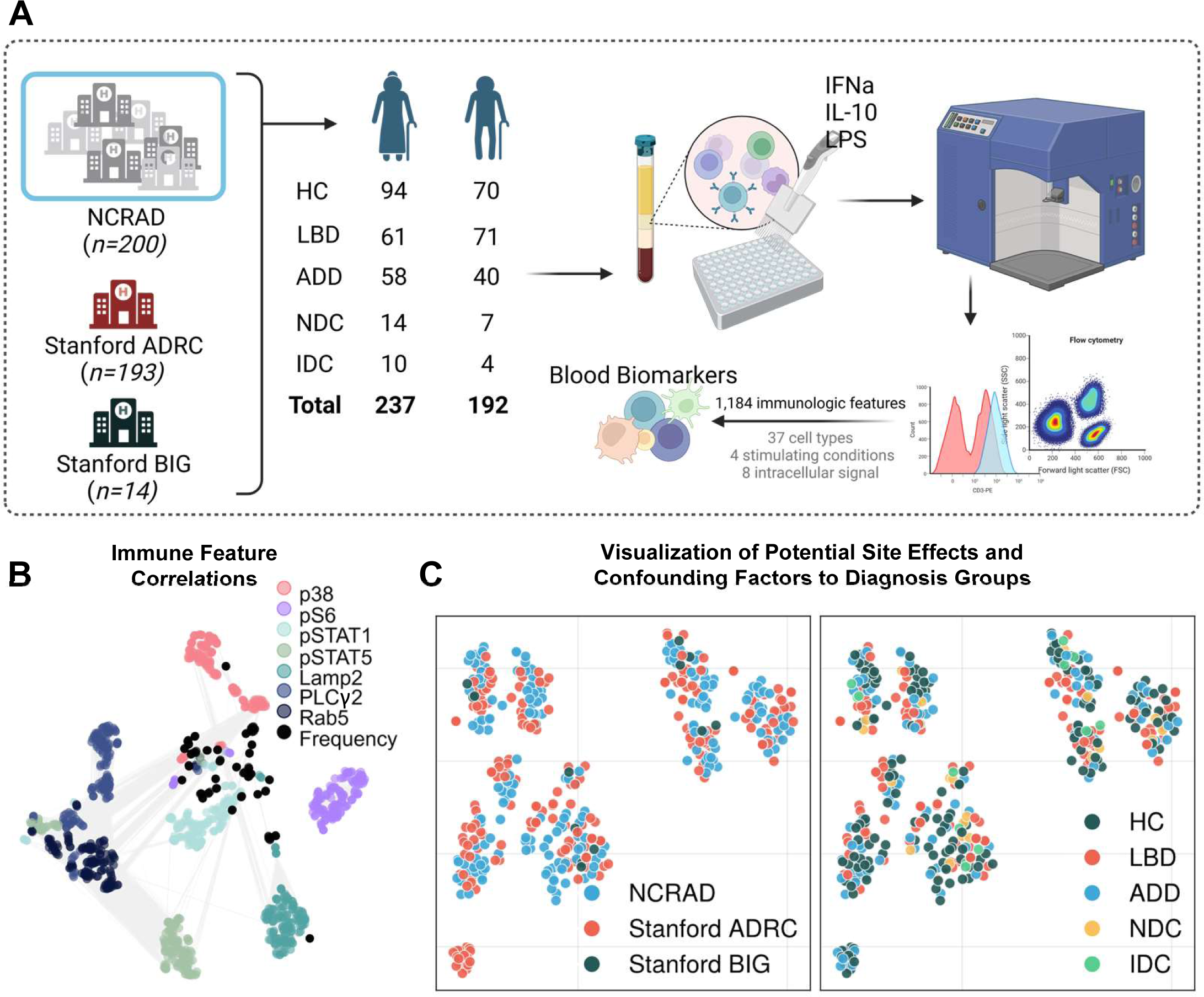
Overall Experiment and Resulting Immune Landscape. **(A)** Diagram of the experiment. PBMCs were collected from diagnosis groups at Stanford ADRC, Stanford BIG, and NCRAD, which in itself aggregated samples from multiple sites. This was followed by stimulating the PBMCs with one of three different canonical immune activators or vehicle control, immunolabelling for surface and intracellular markers, and measuring the cell-specific signals using flow cytometry. Single-cell signals were manually gated to different cell types, resulting in 1,184 immune features for each PBMC sample that were then used by machine learning for the identification of biomarkers. **(B)** A correlation network (edges represent Pearson’s R > 0.7) indicates that the immune landscape was mostly determined by the intracellular signals, i.e. the same intracellular signals tend to be correlated to each other despite different cell types and stimulating conditions. **(C)** The t-SNE plots suggest that there was not a strong effect by the site of sample collection (left), and that samples from different diagnosis groups were well distributed overall (right).

The immune feature landscape (**Fig. 1B**) indicates that, regardless of stimulation and cell type, features from the same intracellular signals tended to be highly correlated with each other, aligning with known intracellular signaling cascades. A subset of pSTAT1, pSTAT5, and pPLCγ2 were highly correlated, whereas pS6 was the least correlated to other signals. A t-SNE plot for patient landscape colored by site indicated that batch correction was effective as there was no apparent site-specific cluster (**Fig. 1C** left). While there could be other confounding factors other than sites, **Fig. 1C** shows that all diagnosis groups were well distributed, hence allaying concerns of any strong effects introduced by confounders.

### Immune Features Differentiate LBD from HC and Other Diseases

The machine learning model (LGBM) exhibited strong performance for separating LBD from HC (Area Under the Receiver Operating Curve [AUROC]=0.90±0.06, Area Under Precision-Recall Curve [AUPRC]=0.86±0.06; **Fig. 2A**), while predictions were essentially random for HC vs. ADD (AUROC=0.52±0.06, AUPRC=0.42±0.05). It should be noted that random guess would yield an AUROC of 0.50, and an AUPRC equivalent to the prevalence of the positive class, which is displayed as patterned gray bars in all figures. The uneven distribution of LBD among sites could be concerning; however, even if the training and test set were split by site, instead of random cross-validation, or if only the Stanford cohort was included, the model still achieved high performance for HC vs. LBD (AUROC=0.77 in **Fig. 2B**; AUROC=0.82±0.08 in **Fig. S2**). This indicates that there was a generalizable pattern of PBMC response for participants with LBD regardless of clinical subgrouping. To ensure that these immune features were unique to LBD, the same HC vs. LBD model was used to predict ADD vs. LBD, NDC vs. LBD, and IDC vs. LBD without retraining. All of these comparisons resulted in high performance with all AUROC above 0.85 (**Fig. 2C**). Corresponding to these AUROC performances, **Fig. 2D** shows that the predicted values for LBD in the test set were significantly different from all other diagnoses. Moreover, the residual of the model predictions (**Fig. 2E**) was not significantly correlated with sex, age, *APOE* epsilon 4 allele status, Levodopa dosage, or subgroup diagnosis of PD vs. PDD; however, the model’s residuals were significantly correlated with DLB vs. PD/PDD, indicating that the model performed equally well across these major variables except diagnosis group DLB compared to PD/PDD.

**Figure 2.**
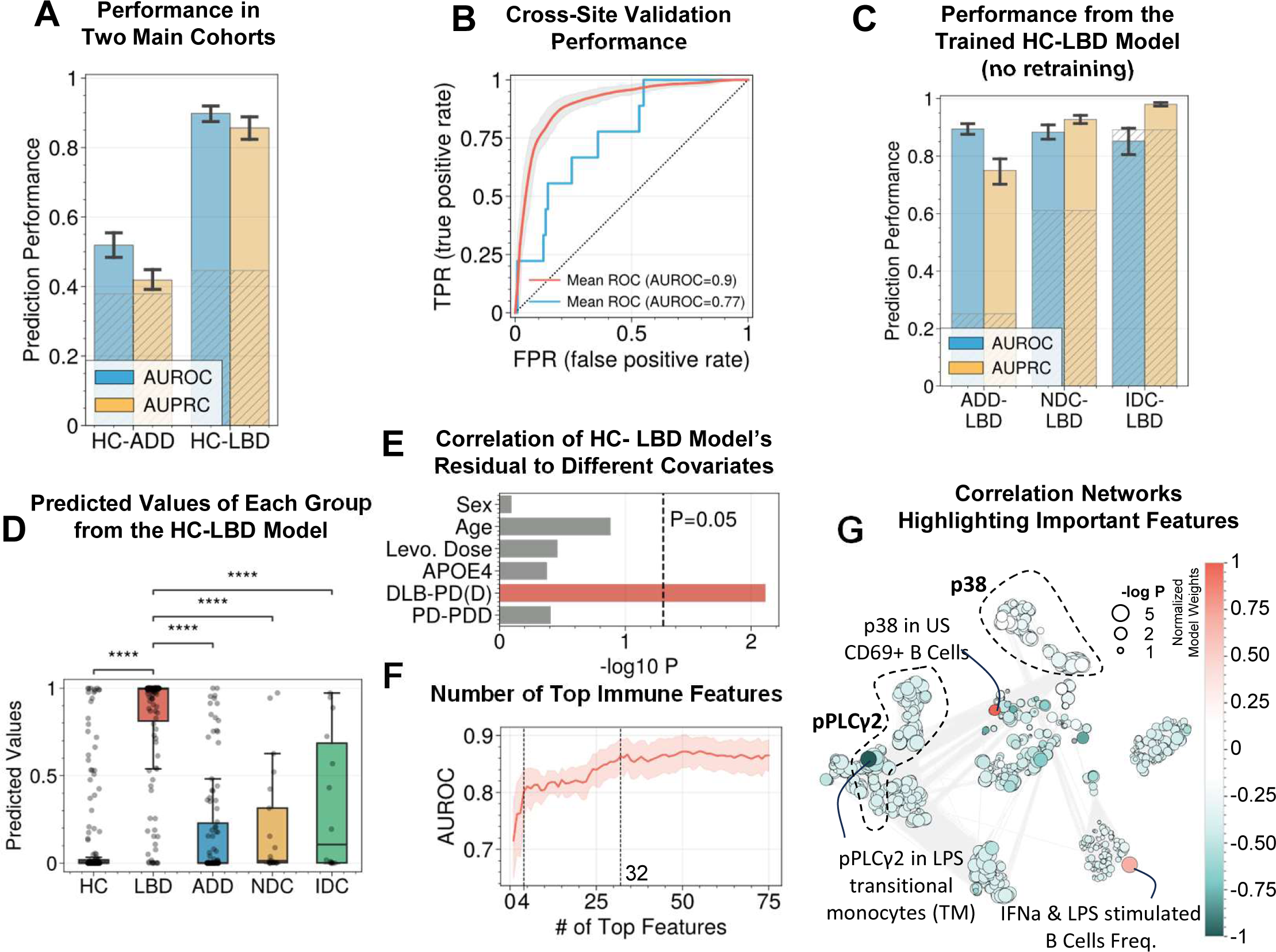
Models developed from multi-site data suggest peripheral biomarkers for LBD. **(A)** The model performance suggested good separation for HC vs. LBD, but not for HC vs. ADD. Note that a random guess baseline would yield an AUROC of 0.50 and an AUPRC equivalent to the prevalence of the positive class in the sample group, which are shown as patterned gray bars. **(B)** Performance using cross-site splitting instead of random cross-validation suggests the generalizability of the biomarkers. **(C)** Transferring the HC vs. LBD model (without retraining) to classify LBD from disease controls, including ADD, NDC, and IDC, yielded similarly high performance. **(D)** The predicted values from the HC vs. LBD model for all diagnosis groups show that the model is LBD-specific. **(E)** Model residual (errors from each prediction) did not significantly (M.W.U. P<0.05) vary with sex, age, Levodopa dosage, APOE e4 status, or PD vs. PDD. This indicates that the model performed equally well across these variables. In contrast, the model’s residual varied for the DLB vs. PD/PDD group, suggesting that the performance of the DLB group differed from the PD/PDD group. **(F)** The required number of top immune features needed to achieve similar performance as all 1,184 features. **(G)** Correlation network highlighting the top features and the immune features with which they are correlated.

Model reduction indicated that only the top 4 immune features were necessary to achieve a satisfactory prediction performance, and 32 features would yield similar performance as using all 1,184 immune features (**Fig. 2F**). The top 4 immune features for LBD were highlighted in the immune feature correlation network (**Fig. 2G**). They include reduced pPLCγ2 response from LPS-stimulated CD14+ CD16+ monocytes, elevated p38 response from unstimulated CD69+ B cells, and frequency of IFNa and LPS-stimulated B cells.

Due to the high correlations among immune features, the model may only select a few representative ones, and interpretation from the model alone may leave out other important biological features. For this reason, other immune features were investigated from a univariate perspective. Heatmaps of the correlations between the top intracellular signals and LBD diagnosis show cell type-specific signals, including: reduced expression of pPLCγ2 in CD69+ NK cells, transitional monocytes (TM), and CD11b+HLA-DR+ TM; reduced expression of pSTAT5 in multiple CD4+ cells; and elevated expression of p38 in multiple CD4+ and CD8+ cells in patients with LBD compared to HC (**Fig. 3A**). Notably, these signals were significantly different between LBD vs. HC and LBD vs. ADD but not between LBD vs. NDC or LBD vs. IDC (**Fig. 3B**), highlighting the needs to integrate multiple immune features and non-linear models.

**Figure 3.**
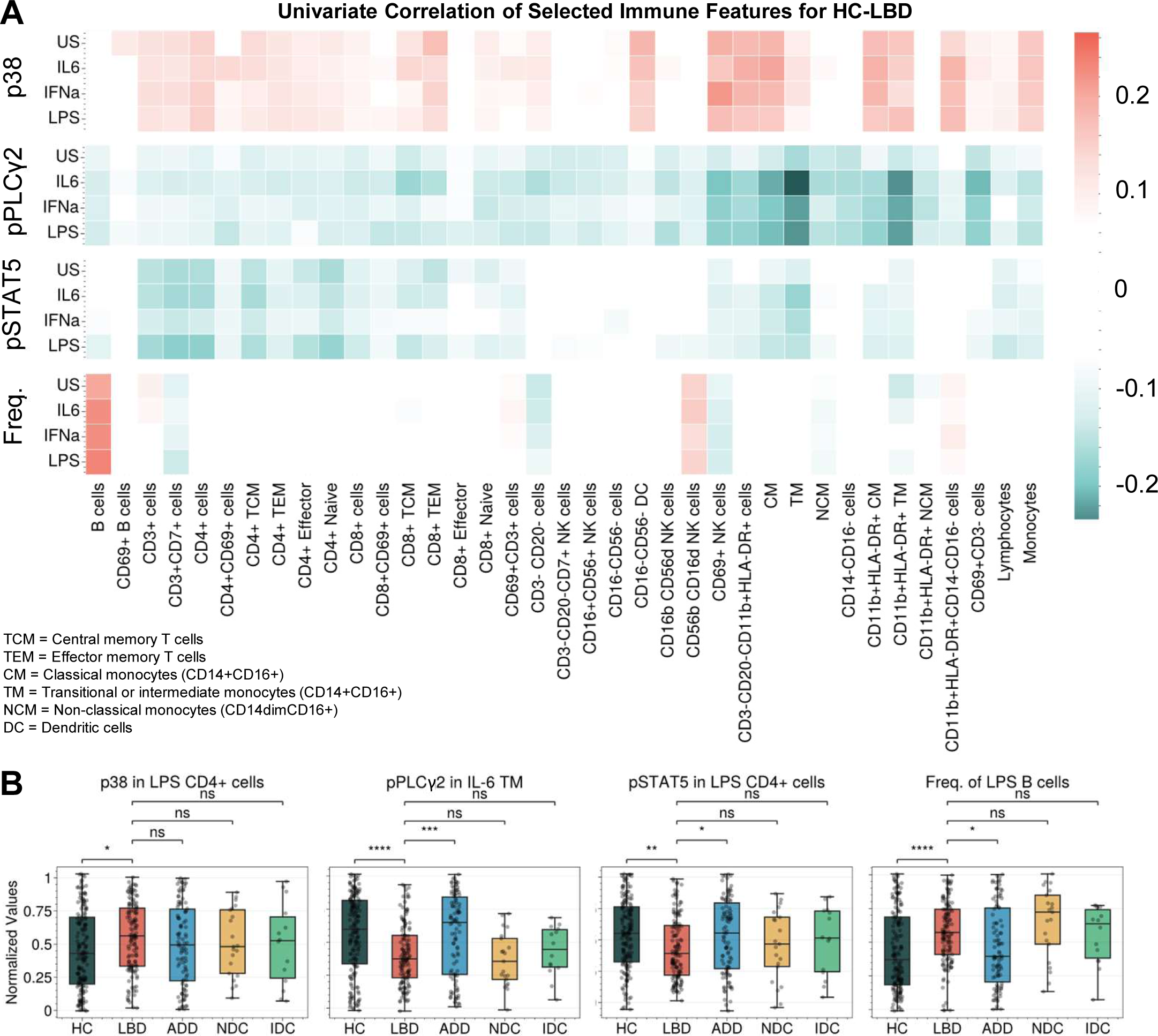
Strong signals for HC vs. LBD were cell-type specific. **(A)** The heatmap of selected intracellular signals (or frequency) from all cell types shows the cell types with the strongest correlations to LBD. **(B)** Examples of the top univariate immune features.

### Differential Signals Separating DLB, PD, and PD with Cognitive Impairment

So far, we have determined a unique peripheral immune pattern for patients with LBD compared to HC, ADD, and other neurodegenerative or autoimmune disease controls. However, as noted above, patients with LBD are a mix of individuals with three different clinical diagnoses (PD, PDD, and DLB) that can be difficult to distinguish clinically with precision and that can merge over time. Our results show that each of these diagnostic subgroups of LBD can be separated from HC moderately well with HC vs. DLB exhibiting the lowest performance (AUROC=0.70-0.93, AUPRC=0.35-0.87; **Fig. 4A**). Transferring these models without retraining to cross-predict among themselves, *e.g.* PD vs. PDD or PDD vs. DLB, exhibited moderately low performance (AUROC=0.60-0.71; **Fig. 4B**). The moderate classification performance indicates that PD, PDD, and DLB share some critical PBMC immune responses in addition to the known shared neuropathological features. Interestingly, the model transfer to classify each LBD subgroup vs. ADD resulted in high AUROC (>0.89) for both PD and PDD (**Fig. 4B**) but not as high for ADD vs. DLB (AUROC=0.67), perhaps because of the well-described comorbidity between DLB and AD neurodegenerative change in the majority of people diagnosed clinically with DLB^37^.

**Figure 4.**
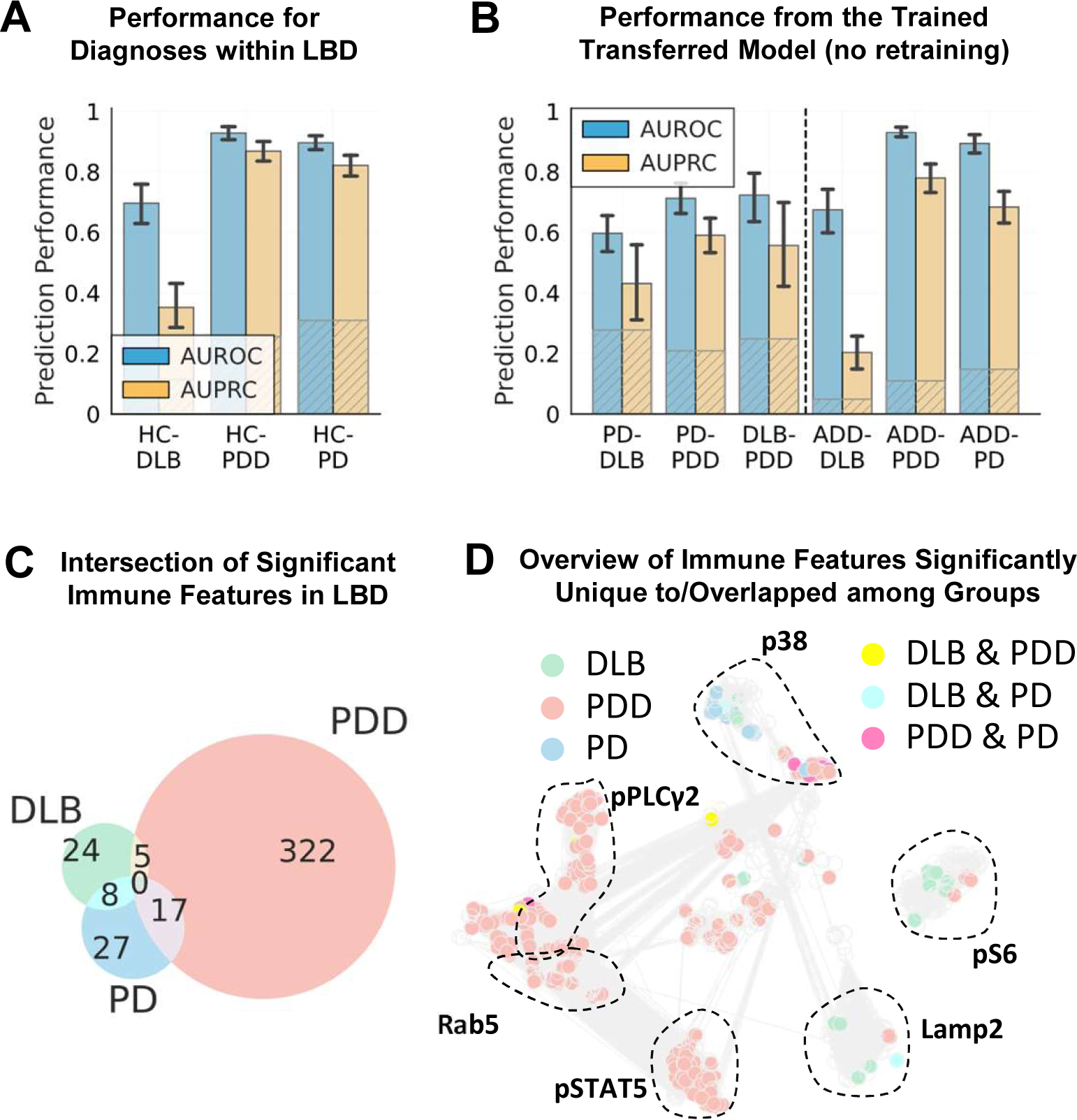
All subgroups within LBD can be separated from HC, but not among themselves. **(A)** Model performance of three separate models each developed for classifying HC from each of the subgroups within LBD, including DLB, PDD, and PD. **(B)** The performance of the same models (without retraining) classifying among each of the subgroups and all of them vs. AD. **(C)** The Venn diagrams of significant immune features for each group (M.W.U. P<0.01) indicated small overlapping features among them. **(D)** The correlation network shows which immune features were unique to or overlapping between DLB, PDD, and PD.

From a univariate perspective when compared with HC, PDD exhibited the highest number of statistically significant immune features (M.W.U. P<0.01), and only a handful of these was shared by PD and DLB (**Fig. 4C**). From the univariate intracellular signals for LBD in the previous section, elevated p38 responses were uniquely associated with a diagnosis of PD with or without dementia (**Fig. 4D**), while most of the reduced pPLCγ2 response and reduced expression of pSTAT5 were uniquely associated with PDD only.

Cognitive exams in multiple domains are predictive of cognitive status in LBD^38^, and together with motor exams and clinicians’ judgment, were the source for deriving a clinical diagnosis. We also tested if the immune features can predict any of the 18 neuropsychological battery test scores in the cases where the data were available, such as trail making or MMSE, or any of the 23 motor examinations from the Unified Parkinson’s Disease Rating Scale (UPDRS) among patients with LBD. Our results show moderately low performance, indicating that the selected immune features were not specific to these measurements in LBD patients (**Fig. S3 & S4**).

### The Biological Pathways from the Identified Biomarkers did not Overlap with Other Comorbidities

Several diseases and conditions that are not primarily associated with neurodegeneration tend to increase or lower the risk of dementia and PD. Examples of these include arthritis^39^, diabetes^40^, hypercholesterolemia^41^, hypertension^42^, REM sleep disorder^43^, sleep apnea^44^, traumatic brain injury (TBI)^45^, and vitamin B12 deficiency (VB12DEF)^46^. This section aims to investigate whether the peripheral immune biomarkers discovered above had links with these common comorbidities. In the cases where comorbidities data were available in our sample set, individuals with these comorbidities were almost equally split among HC, ADD, or LBD (**Fig 5A**) except for diabetes, which only occurred in HC and AD, and REM sleep disorders, which only occurred in LBD. Models developed to test if the collected PBMC immune features were able to predict these comorbidities showed that none of the comorbidities could be predicted accurately with the marker panel selected for this study (**Fig. 5B**). Indeed, only TBI and VB12DEF achieved AUROC above 0.60. This is further supported by a univariate analysis showing that there was minimal overlap of significant features (M.W.U. P<0.01) between TBI, VB12DEF, and LBD (**Fig. 5C**). Together, these results suggest that the biomarkers identified were unique to LBD and were minimally influenced, if at all, by these comorbidities.

**Figure 5.**
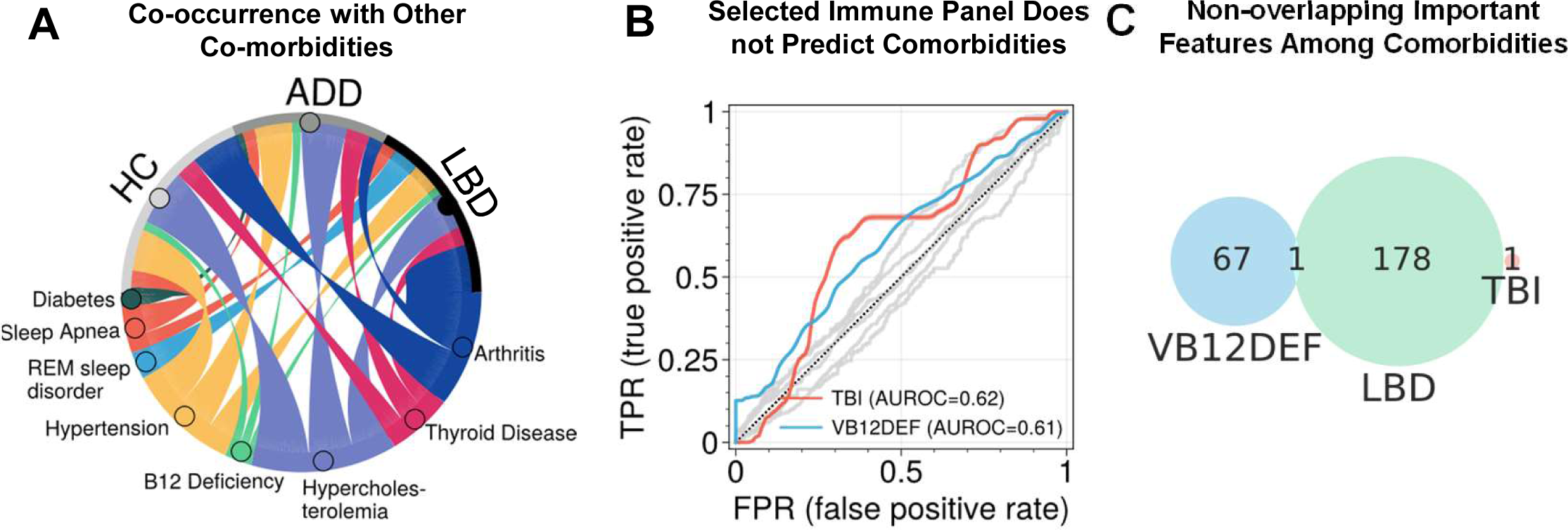
The identified LBD biomarkers did not have overlapping biological pathways with common non-neurodegenerative comorbidities. **(A)** A chord diagram displaying LBD, ADD, or HC co-occurrence with other comorbidities. Note that TBI was also included but due to a low number of cases (*n=*6), it is now shown in the plot. **(B)** Model performances (AUROC) for all comorbidities were below 0.60 except for TBI and vitamin B12 deficiency (VB12DEF). **(C)** The Venn diagrams of significant immune features for each group (M.W.U. P<0.01) indicated no overlapping features among them.

## Discussion

Human genetic, pathologic, imaging, and biochemical data as well as results from experimental models have linked neuroinflammation with the initiation or progression of prevalent age-related neurodegenerative diseases. Among these, the LBD spectrum, PD, PDD, and DLB, have been most strongly linked to events in the periphery as potential contributing mechanisms that impact the brain^47^. Here we tested the hypothesis that cell-specific immune responses by PBMCs might be associated with LBD diagnosis, highlighting potential peripheral biomarkers and possibly illuminating mechanisms of disease. Our multisite study design included PBMCs from 429 participants from five diagnostic groups (HC, LBD, ADD, and NDC as controls for non-specific changes occurring with debilitation from neurodegenerative diseases, and IDC to control for non-specific changes occurring with immune-mediated diseases) that were investigated in basal state or following stimulation by canonical immune activators to generate 1,184 molecular features per individual. These rich immune response data were coupled with extensive clinical annotation and analyzed by machine learning techniques.

Our major finding was that, within the context of our stimulants and multiplex panel, only 4 immune features were necessary to achieve similar prediction performance for LBD as all immune features; these were: reduced pPLCγ2 response from LPS-stimulated transitional monocytes, elevated p38 response from unstimulated CD69+ B cells, and increased frequency of IFNa and LPS stimulated B cells. Together these data suggest a broad alteration in peripheral immune response in patients with LBD that is distinct from other neurodegenerative and autoimmune diseases, and that involves monocytes and lymphocytes. Although these findings establish relevance to the human condition, determining the mechanisms by which these stimulant- and cell-specific immune responses may or may not directly contribute to LBD-type neurodegeneration will require means of selectively manipulating each in isolation or combinations in model systems that faithfully reflect the human immune system and mechanisms of neurodegeneration in LBD.

On top of identified features from the model, univariate statistical analysis results highlight three immune response features that are strongly characteristic of PBMCs from people diagnosed with LBD: reduced pSTAT5 in CD4+ subset and reduced pPLCγ2 response and elevated p38 response in subsets of NK cells and TM cells. Our localization of elevated p38 response to lymphocytes in people with LBD suggests that this may be a feature of a subset of lymphocytes that traffic into the brain as immune master regulators^47^. Additionally, p38 is extensively related to gut immunity, inflammation, and aging^48–50^; gut physiology has been implicated by many studies as a potential contributor to LBD^51^. PLCg2 is highly expressed in immune cells including microglia, and gain-of-function mutations in *PLCG2* cause autoimmune diseases^52–55^. A nonsynonymous variant in *PLCG2* is associated with reduced risk of ADD, DLB, and frontotemporal dementia, suggesting a broad influence on the mechanisms of neurodegeneration, most likely neuroinflammation^33,56^. Our results showed reduced phosphorylation of PLCg2, the molecular mechanism of its activation, in peripheral monocytes and other PBMCs of patients with LBD, thereby aligning with genetic data associating less active PLCg2 with increased risk of LBD. In a previous single-site study we identified reduced pPLCγ2 in a small group of ADD participants^25^; however this result did not generalize to the current multisite study with 4 times more ADD samples. Together, these findings suggest a broad influence of PLCg2 activation in peripheral immunocompetent cells in multiple forms of neurodegenerative disease but most robustly in LBD.

The medical and pathological distinctiveness of the LBD subgroups, PD, PDD, and LBD, is a decades-long debate^5^. We sought to determine the extent to which peripheral immune responses as measured here may potentially point to LBD subgroup-specific features. We observed low model prediction performance among PD, PDD, and DLB suggesting that at least as determined by our multiplex panel, PBMC immune responses are similar among the three subgroups. Further univariate analysis suggested that increased signaling through pPLCγ2 and pSTAT might be a peripheral immune feature specific to PDD and not PD or DLB. Interestingly, despite being predictive of LBD and its subgroups, peripheral immune responses were not strongly predictive of performance on neuropsychological tests or consensus motor evaluation, nor were they associated with other medical conditions shown to modulate the risk of LBD. We speculate that the detected peripheral immune response in LBD subgroups may be a consequence of LBD-type neurodegeneration or may reveal an underlying inherited or acquired trait that renders a person more vulnerable to developing LBD without being directly involved in the extent of neurodegeneration.

Our study has limitations. While the overall sample size is adequate, some of the LBD subgroup sizes were small and lacked neuroimaging, biomarkers, or pathologic validation of clinical diagnosis. For these reasons, LBD subgroup comparisons should be considered preliminary. Also, the multisite samples were majority Caucasian or Asian representing a national deficit in sample diversity among these diseases that is currently being addressed. With these limitations in mind, our quantification of PBMC immune response from multisite research participants yielded a unique pattern for LBD compared to HC, multiple related neurodegenerative diseases, and autoimmune diseases thereby highlighting potential biomarkers and insights into mechanisms of LBD.

## Acknowledgments

Samples from the National Centralized Repository for Alzheimer’s Disease and Related Dementias (NCRAD), which receives government support under a cooperative agreement grant (U24 AG21886) awarded by the National Institute on Aging (NIA), were used in this study. We thank contributors who collected samples used in this study, as well as patients and their families, whose help and participation made this work possible. Support for this research related to healthy controls, ADD, and LBD samples has also been kindly provided by the Stanford ADRC and the Pacific UDALL Center. Support for this research related to autoimmune disease controls has been kindly provided by the Project BIG Fund and the Garrett Immunology Research Fund. Flow cytometry and initial analysis thereof were performed in the Stanford Human Immune Monitoring Center (HIMC).

This work was supported by NIA RF1AG077443 (T.J.M., N.A.), NIGMS 5T32GM089626 (P.C.). Funding also provided for by the Alzheimer’s Drug Discovery Foundation (ADDF) Diagnostics Accelerator, a fund set up in collaboration with Bill Gates and other philanthropic partners. The initiative seeks to accelerate the development of affordable and accessible biomarkers to diagnose Alzheimer’s disease and related dementias, and to advance the development of more targeted treatments. To learn more about the initiative visit the website at: (AlzDiscovery.org).

Conceptualization, T.P. and T.J.M.; Methodology, T.P., C.R.G., K.M., H.T.M., F.X., C-H.S., N.A., and T.M.; Investigation, T.P., N.S., K.M., E.J.F., S.C.P., and S.Y.W.; Visualization, T.P., C.E., A.P.; Writing – Original Draft, T.P. and T.J.M.; Writing – Review & Editing, E.B., D.H., B.C., M.R., P.C., N.A., and K.S.M.; Data Curation, D.C., L-J.H., T.P., K.M.; Funding Acquisition, T.P., E.J.F., T.J.M., N.A., K.L.M., K.S.M.; Resources, D.C., L-J.H., J.E.D., and L.B.K.; Supervision, T.P., N.A., and T.J.M.

## Methods

### Study Design

This study aimed to determine whether differences in peripheral immune responses between healthy controls (HC) and research participants with LBD (PD, PDD, and DLB) are detectable by flow cytometry analysis of PBMCs. In addition, we included samples from other research participants for neurodegenerative disease controls (NDC) and patients with autoimmune diseases for immune disease controls (IDC) to control for nonspecific effects of debilitation from neurodegeneration and immune-mediated diseases. Participants were research volunteers at Stanford Alzheimer’s Disease Research Center or the Pacific Udall Center (Stanford ADRC), Stanford BIG Project (BIG), and many other Alzheimer’s Disease Research Centers (ADRCs), whose samples were aggregated and distributed by the National Centralized Repository for Alzheimer’s Disease and Related Dementias (NCRAD). All participants provided written informed consent to participate in the study, which followed protocols approved by the Stanford Institutional Review Board. Clinical diagnosis was made by consensus criteria.

Blood was collected from a total of 429 volunteers stratified into seven diagnosis groups: HC (*n*=164), LBD (total n=132 including 67 PD without dementia, 47 PD with dementia (PDD), and 18 DLB), Alzheimer’s disease dementia (ADD, *n*=98), other neurodegenerative disease controls (NDC; *n*=21), and immune disease controls (IDC; *n*=14). The diseases included in NDC were multiple system atrophy, primary supranuclear palsy, corticobasal degeneration, frontotemporal lobar degeneration, behavioral frontotemporal dementia, primary progressive aphasia, vascular brain injury, prion disease, and traumatic brain injury. HCs were individuals who were not diagnosed with any neurological disease and had no cognitive impairment. AD, LBD (PD/PDD/DLB), and NDC participants had a single clinical diagnosis without clinical comorbidity. The sex distribution of each group is shown in Fig. 1A, and the average age was 73±6 for HC, 75±8 for AD, 71±7 for PD, 73±7 for PDD, 73±7 for DLB, 74±7 for NDC, and 67±3 for IDC. The race distribution of participants who contributed to our sample set was 86% White, 12% Asian, 1% Black or African American, and 1% Others. The percent contribution of each diagnosis group from each site was 35% Stanford ADRC and 65% NCRAD for HC, 36% Stanford ADRC and 64% NCRAD for ADD, 93% Stanford ADRC and 7% NCRAD for LBD, 100% NCRAD for NDC, 100% BIG for IDC. The protocol for PBMC collection and storage by each site can be found in the **Supplementary Materials**.

### Flow Cytometry Experiment

PBMCs were isolated by density-gradient centrifugation and cryopreserved. Post-thaw, cells were washed in a complete RPMI medium with benzonase. Cell viability as measured by Vi-Cell (Beckman Coulter) for all samples was above 90%. After resting for 2h at 37oC, PBMCs were either left unstimulated or stimulated with a panel of cytokines: IFNα (10,000 units/ml), IL-2 (50 ng/ml), IL-6 (50 ng/ml) and LPS (200 ng/ml) for 15 min, at 37oC. Stimulation was stopped by fixing cells with paraformaldehyde for 10 minutes at room temperature. After washing cells with PBS, samples were stained with LIVE/DEAD™ Fixable Blue Dead Cell Stain Kit, for UV excitation (from Invitrogen) for 15 min at room temperature. After live dead staining, cells were washed with wash buffer (Phosphate buffered saline, 2% Fetal bovine serum, 0.1% sodium azide), followed by surface staining with anti-CD4(BUV805), CD7 (AF780), CD8 (AF700), CD11b (BUV395), CD14 (BUV737), CD16 (BV750), CD19 (PerCP-Cy5.5), CD27 (BV711), CD56 (BUV563), CD69 (BUV661), HLA-DR (BV480) (antibodies from BD Biosciences), CD3 (BV605) and CD45RA (BV570) (antibodies from BioLegend). Staining was done at room temperature for 30 min. After 2 washes, cells were permeabilized with ice-cold methanol and were stored overnight at −80°C. Post permeabilization, cells were washed again, and intracellular staining was done with anti-pSTAT1 (AF488), pSTAT5 (PE-Cy7), pP38 (PE), pPLCγ2 (APC), pS6 (BV421), CD107b/Lamp2 (BV786), (antibodies from BD Biosciences) and Rab5 (PE-CF594) (from Santa Cruz Biotechnology) at room temperature for 30 min. After two further washes, the acquisition was performed on a BD Symphony A5 flow cytometer with a High Throughput Sampler (HTS) and analyzed using FlowJo software where median expressions were collected for each gated cell type. The reagents and the gating scheme can be found in **Table S1** and **Fig. S1**. Lastly, Combat^57^ was used to correct site effects.

### Data Analysis

Machine learning is a common tool for extracting insight from high-dimensional cytometry data^58,59^. Here, light gradient-boosting machine (LGBM)^60^ was used as it outperformed other machine learning models, including logistic linear, random forest, and feed-forward neural network models, in our dataset. To maximize generalizability, the performance was evaluated using 10 repeated 4-fold cross-validation where the model is trained on a randomized train set and tested on unseen samples. For the classification of the three main groups, HC and IDC were merged and labeled 0, and the disease group (LBD or ADD) was labeled 1. The test set prediction values were used for subsequent analyses and visualizations. The model performance metrics include the Area Under the Receiving Operating Curve (AUROC) and the Area Under the Precision-Recall Curve (AUPRC). For differential predictions, *e.g.* ADD vs. LBD or NDC vs. LBD, the primary model trained for HC vs. LBD was used without retraining. For the prediction of LBD subgroups (PD, PDD, DLB), comorbidities, and motor examinations, LGBM was also used with the same cross-validation setup except that a subsampling technique was used to ensure balanced age and sex ratios between case and controls. Methods for model reduction and correlation networks can be found in the **Supplementary Materials**.

## Supplemental Information

Data availability: Singlet live cell data (.fcs format), gated median value data (.csv format), the associated metadata (.csv format), and the data dictionary are made publicly available at DOI: https://datadryad.org/stash/share/LT4qx1N_pGC5WlOo24QNDt7R61BglFXnxuK7qgjvTpE.

