## Supplementary Materials for "Single-Cell Peripheral Immunoprofiling of Lewy Body Disease in a Multi-site Cohort"

**PBMC Collection and Storage at Stanford ADRC**

1. Pour blood into one 50ml conical tube (should fill up to around 40-45 ml)

a. Save one label for later to put in yellow notebook.

2. Centrifuge with balance for 10 mins at 3000 rpm (check brakes are 5 for acceleration and deceleration. Assure that the temperature is 20 degrees C.)

3. If you disturb the blood beneath plasma, spin again at 3000 rpm for 3-4 mins.

4. Take off the plasma from the top and make 12- 1ml and 0.5 ml tubes (labeled with PIDN, year, P for plasma, volume) on tops and sides. Write your initials on side of tubes.

5. Shake then add PBS equal to blood volume to the 50ml conical tube of blood (Max volume = 40ml). Mix well with electrified pipette.

6. Measure out 15 ml of Ficoll into one new 50ml conical tube for low blood volume or two 50ml conical tubes for high blood volume (above 30ml).

7. Very slowly add the PBS+Blood mixture into the Ficoll Tube(s) (10 ml at a time). There should be 20 ml added to each. Make sure it forms a layer on the top of the Ficoll. Balance tubes.

8. Centrifuge PBS+Blood mixture and Ficoll for 30 minutes at 1500 rpm.

a. During the break, put away plasma, add samples to box map, and label 10 PBMC tubes with PIDN, Year, “PBMC”, and date (screw cap tubes)

9. Take out the tiny cloud layer with a plastic pipette from both tubes and put in a new 50ml conical tube.

10. Shake then add PBS up to 40ml and spin for 10 minutes at 1500 rpm

a. Remove Soultions A and B from 4-degree fridge and allow to acclimate to room temp.

11. Carefully pour off PBS from the pellet into one of the empty tubes and toss.

12. Add PBS to 40 ml again and tap the pellet to mix it.

13. Spin again for 10 minutes at 1500 rpm

14. Pour off PBS carefully (don’t pour off pellet), use a Kimwipe to absorb the extra.

15. Add 1ml of Solution A to the pellet and mix by pipetting up and down.

16. Take the tiny plasma tube and add 10 ul to 10 ul of Trypan blue and mix up and down.

17. Add 10 ul to the hemocytometer.

18. Count the first 4 boxes using the microscope and add up the # of cells.

a. Multiply sum by 8 and add 4 zeros to get the number of cells in millions

b. Divide by 5,000,000 to get the number of tubes you would need (each tube should have around 5 million cells per 0.5ml)

19. Divide # of tubes needed by 2 and that is the total ml of solution you need to make with equal parts solution A and solution B (factor in that 1ml of Solution A was added).

20. Add 0.5 ml to each labeled 1.2ml cryovial tube and add to Mr. Frosty

21. Add Mr. Frosty to the -80 freezer.

22. Complete sample records in Excel sheets “ADRC - Plasma and ADRC-PBMC” (be sure to record fasting and time of draw from clinical sheet) and RedCap.

**PBMC Collection and Storage at NCRAD**

1. PBMC Isolation

1.1. Centrifuge NaHep tubes at 800 x g for 10 minutes to isolate the buffy coats.

1.2. Using a sterile alcohol prep pad, carefully remove the green rubber stopper from each tube

and place on an ethanol-soaked paper towel.

1.3. Using a 5 mL serological pipette, remove the top layer of plasma, leaving about ½ inch of

plasma on top of the buffy coat.

1.4. Gently collect the buffy coat layer in a slow, circular motion (~2 mL per tube).

1.4.1. It is ideal to collect even amounts of plasma and RBCs with the buffy coat collection.

1.4.2. Turning down the speed of the Pipette Aid may help in collecting all of the buffy coat.

1.4.3. It is easier when processing several samples at a time to collect even amounts of buffy coats across each sample. Aim to collect exactly 2 mL per blood tube.

1.5. Take note of the volume of the buffy coat. Dispense the buffy coat into one 14 mL

polystyrene tube.

1.6. If combining two samples from the same kit, repeat steps 1.2 through 1.5 to combine the

buffy coats together for processing. Do not exceed 6 mL of total buffy coat per one 14 mL

polystyrene tube. Only combine buffy coats from the same kit—DO NOT combine buffy coats

from different kit numbers.

1.7. Add 0.5 M EDTA to each sample. EDTA volumes depend on the total amount of buffy coat

collected per sample. See Appendix D for volumes.

1.8. Add the Isolation Cocktail at 50 µL/mL. Isolation Cocktail volumes depend on the total

amount of buffy coat collected per sample. See Appendix D for volumes.

1.9. Mix by using a 1,000 µL pipette set to 1,000 µL or a 5 mL serological.

1.10. Incubate at room temperature for 5 minutes. This incubation may go a couple minutes over the stated incubation time, if needed.

1.11. Add a 1:1 amount of DPBS and mix 2-3 times by pipetting. (If 2 mL of buffy coat was

collected, then add 2 mL of DPBS.)

1.12. Essential Step: Vortex the RapidSpheres™ for a minimum of 30 seconds directly before

use.

1.13. Add the RapidSpheres™ at 50 µL/mL. RapidSpheres™ volumes depend on the total

amount of buffy coat collected per sample. See Appendix D for volumes.

1.14. Mix by using a 5 mL serological pipette or a 1,000 µL pipette set to 1,000 µL.

1.15. Immediately after mixing the RapidSpheres™, place the sample tubes in the magnet,

leaving the lids OFF.

1.16. Incubate the samples in the magnet for 5 minutes at room temperature. This is the first

magnetic incubation.

1.17. After the incubation, carefully pipette along the opposite side of the tube from the magnet,

transferring the cell suspension into the second 14 mL polystyrene tube.

1.17.1. Note: When collecting the cell suspension, RBCs will also be collected. Collect up to 10% of the starting buffy coat volume (0.2—0.8 mL) of RBCs from the bottom of the sample. Some liquid will be left in the original tube. This is normal and to be expected. (For example, if 2 mL of buffy coat was collected, aim to collect ~0.2 mL of RBCs from the bottom of the collection tube.)

1.18. Add the RapidSpheres™ at 50 µL/mL. RapidSpheres™ volumes depend on the total

amount of buffy coat collected per sample. See Appendix D for volumes.

1.19. Mix by using a 5 mL serological pipette or a 1,000 µL pipette set to 1,000 µL.

1.20. Immediately after mixing the RapidSpheres™, place the sample tubes in the magnet,

leaving the lids OFF.

1.21. Incubate the samples in the magnet for 5 minutes at room temperature. This is the second

magnetic incubation.

1.22. After the incubation, carefully pipette along the opposite side of the tube from the magnet,

transferring all of the clear cell suspension into the last 14 mL polystyrene tube.

1.23. Incubate the samples in the magnet for 5 minutes at room temperature. This is the third and last magnetic incubation.

1.24. After the incubation, carefully pipette along the opposite side of the tube from the magnet,

transferring all of the cell suspension into the 50 mL conical tube.

1.25. Bring the final volume up to 30 mL with DPBS.

1.26. Centrifuge the 50 mL conicals to pellet the cells at 800 x g for 10 minutes.

1.27. Decant and discard the supernatant leaving the cell pellet intact.

1.27.1. If the sample has excess RBC contamination, add 10 mL of pre-warmed HYL buffer and pipette up and down to break up cell clumps. If there is no RBC contamination, continue on to section 5.3.

1.27.2. Incubate the samples at RT for 10 minutes.

1.27.3. Bring volume up to 30 mL with DPBS and invert tube to mix cells.

1.27.4. Centrifuge the 50 mL conicals to pellet the cells. Centrifuge at 800 x g for 10 minutes.

1.27.5. Decant and discard the supernatant leaving the cell pellet intact.

2. Cell Counting (LUNA-FL™)

2.1. Re-suspend the cell pellet in 2 mL of DPBS. Briefly vortex the cell suspension to mix well,

1-2 seconds.

2.2. Immediately pipette 18 µL of the cell solution into a microcentrifuge tube, numbered the

same as the sample tube.

2.3. Proceed to count cells with the LUNA-FL™ Cell Counter following SOP IUGB-3-62 CELL

COUNTING-LUNA FL.

2.3.1. Record Viability and Live cell count/mL in Appendix B or C for use in calculations.

3. Cytoslides

3.1. If the study calls for it, calculate the amount of cells and volume needed for the cytoslide

using Appendix C.

3.2. Load and run the Shandon Cytospin.

3.2.1. Place the slide from 5.1.4.7 in the metal mousetrap holder with the labeled side facing away from the holder.

3.2.2. Place the funnel against the slide.
3.2.3. Check the back of the mousetrap to be sure the hole in the funnel aligns with the

coverslip on the slide.

3.2.4. Add an amount of DPBS to the sample so the total volume is equal to the final number of aliquots (refer to the example below).

3.2.4.1. Example: If the calculations in Appendix C suggest 5 aliquots at 1 mL each, bring the sample volume up to 5 mL by adding 3 mL of DPBS. 2 mL of DPBS is lready in the sample tube for counting from step 5.3.1.

3.2.5. Use this cell suspension to load the calculated volume V2 (from the calculations in Appendix C) into the cytospin funnel.

3.5.1. Add the cell suspension to the elbow of the funnel (see below).

3.2.6. Cap all funnels.

3.2.7. Centrifuge at 1000 rpm for 4 minutes with medium acceleration in the Thermo

Shandon Cytospin Centrifuge.

3.2.7.1. When the spin is finished, remove the funnel from the slide by pulling the funnel straight away from the slide. DO NOT drag or touch the funnel against the slide at all.

4. Final Wash and Aliquoting Cells into Cryovials

4.1. Add DPBS to wash the cell solution (from step 5.4.2.4) until the total reaches 30 mL.

4.2. Mix the cell suspension by inverting the tube 3 times.

4.3. Determine the number of cryovials for the freeze.

4.3.1. Freeze at a concentration of 3.0 X 106 cells/mL according to the calculations in Appendix B or C.

4.4. Locate PBMC working list in the database and/or Excel files.

4.4.1. Create children and print labels. See Appendix A for OnCore data entry fields and

related instructions.

4.4.2. Label the cryovials.

4.4.3. Ensure the children aliquot labels match the parent labels. Depending on the study, ensure the kit numbers, ST numbers, or sequence numbers on the children aliquots match the parent labels.

4.5. Centrifuge the samples from 4.3 at 800 x g for 10 minutes.

4.5.1. Make freeze media during this spin. Prepare the freeze media fresh daily, using 45% RPMI, 45% FBS, and 10% DMSO. Make enough freeze media for each cryovial plus one extra per parent sample.

4.6. Decant and discard the supernatant leaving the cell pellet intact.

4.7. For each cryovial needed, use 1 mL of freeze media to re-suspend the cells.

4.7.1. Example: If the calculations in Appendix B or C suggest 6 aliquots, then use 6 mL of freezing media to resuspend the cells.

4.8. Resuspend the cell pellet with the correct amount of freezing media. Keep on wet ice. Repeat for all remaining samples.

4.9. Briefly pipette the cell solution to create a homogeneous cell suspension before aliquoting.

4.10 Dispense the cell suspension in 1 mL aliquots into the cryovials and place into a Mr. Frosty

or a cryobox (based upon method for freezing).

5. Cryopreserving the Cells

5.1. Cryopreserve cells using the Mr. Frosty or Corning® CoolCell® Freezing System.

5.1.1. Mr. Frosty Method

5.1.1.1. Add 250 mL isopropanol to a Mr. Frosty freeze unit. Place a sponge around the inside walls. Make sure the Mr. Frosty is chilled to 4℃ in refrigerator. Note: Replace isopropanol every fifth use.

5.1.1.2. Place holder into Mr. Frosty.

5.1.1.3. Place cryovials into holder.

5.1.1.4. Place Mr. Frosty into a -80 ºC Freezer for 8 hours or longer.

5.1.1.5. Transfer the vials to liquid nitrogen storage after at least 8 hours (see section 5.7).

5.1.2. Corning® CoolCell® Freezing System

5.1.2.1. Before beginning cryopreservation process, ensure CoolCell® with filler vials have been chilled at 4 ℃ (place in the fridge the day before or a few hours in advance).

5.1.2.2. After all the vials have been aliquoted, transfer them into a pre-chilled CoolCell® freezing container. Make sure any unused slots in the CoolCell are filled with filler tubes.

5.1.2.3. Place the CoolCell in a -80 ℃ freezer overnight, freezing the cells at a rate of -1 ℃ per minute.

5.1.2.3.1. Note: Clear away any frost and debris from the freezer shelf to ensure proper air circulation around the freezing container. Failure to do so will cause the vials to freeze unevenly.

5.1.2.4. Store vials in appropriate locations in liquid nitrogen after at least 8 hours.

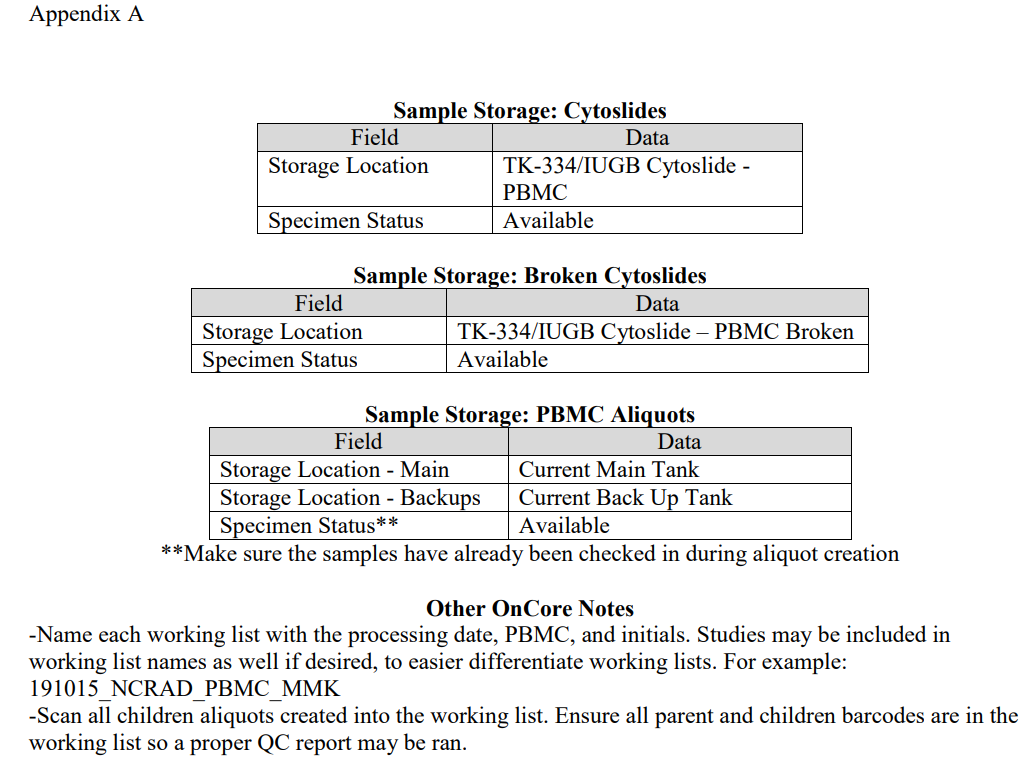

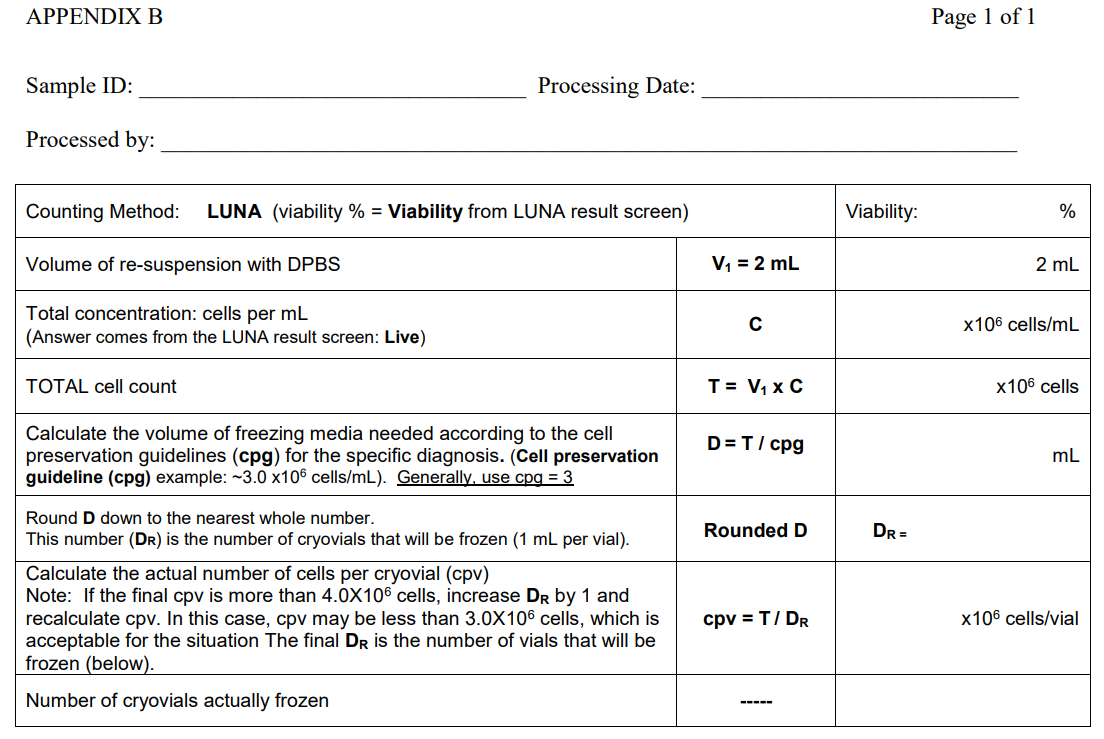

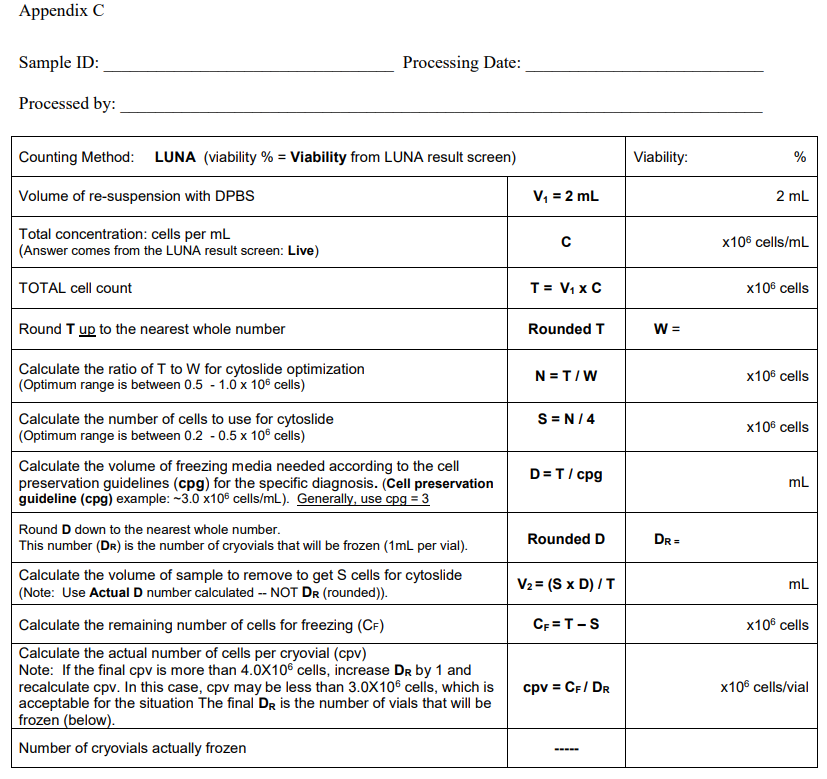

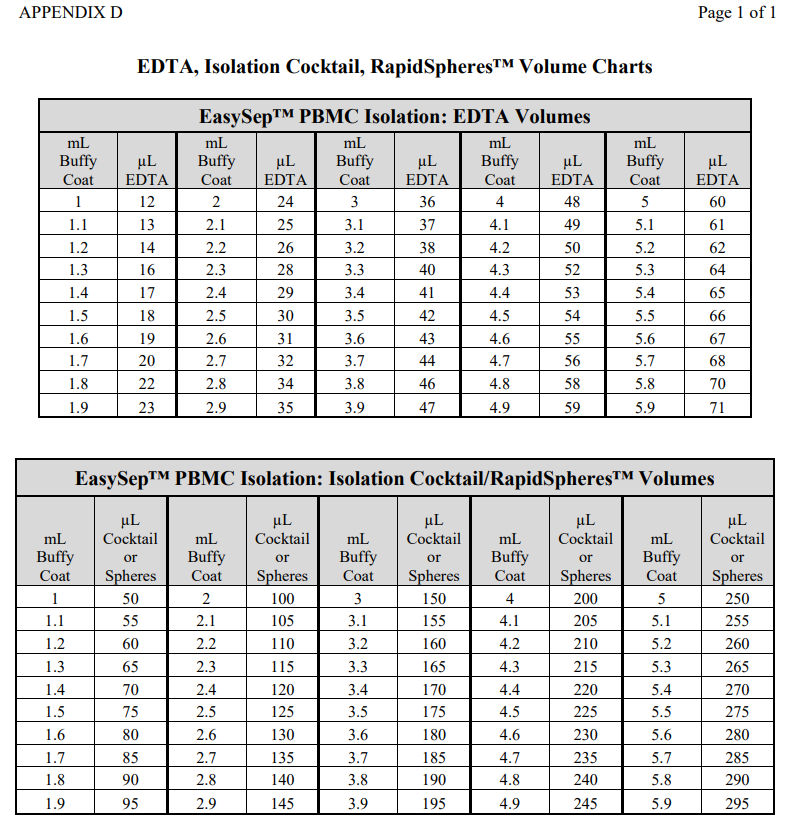

**PBMC Collection and Storage at Stanford BIG Project**

1. Pipette 15mL of Ficoll into the central hole of the SepMate tube.

3. In a 50mL conical tube, measure the volume of heparinized whole blood and add an equal volume of PBS

4. Add the blood to the SepMate tube by pipetting it down the side of the tube a. Add no more than 34mL of blood (no more than 17mL whole blood)

5. Centrifuge the vial at 1,200 x g for 10 minutes with the brake on.

6. Invert the tube (for no longer than 2 seconds) and pour the plasma and PBMCs into a new 50mL conical vial.

7. Add PBS to the tube up to the 50mL mark

8. Centrifuge the PBMCs at 250 x g for 10 minutes.

9. Aspirate the supernatant and resuspend the cells in 48mL of PBS 8 hours.

10. Count the cells using the Tali Counter (or lab’s preferred cell counting method)

10.1. Add 25µL of the cell suspension to a Tali slide

10.2. Choose the “Quick Count” selection and “Name Now”

10.3. Label the data with the sample ID

10.4. Insert slide into the Tali following the arrows on the slide

10.5. Press the button “Press to Insert New Sample”

10.6. Focus the image so that the cells can be seen clearly with definitive borders

10.7. Press “Press to Run Sample”

10.8. After counting, set the cell size to “5µm to 15µm” (this only has to be done the first sample of the day)

10.9. Calculate the total cell count by multiplying the number of cells/mL by the total volume of cell suspension

10.9.1. ex – 3.45 x 105 cells/mL x 48mL =165.6 x 105 cells

11. Centrifuge the conical vial at 250 x g for 10 minutes.

a. Based off the total cell count, calculate the number of vials and volume of freezing media that will be needed (see Table below)

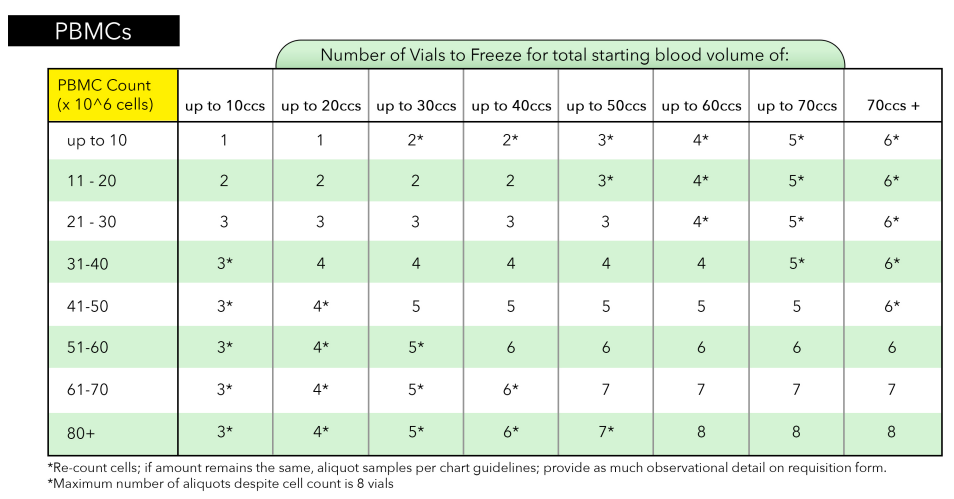

11.1. Label the appropriate number of empty cryovials with deidentified cryogenic label and place in a CoolBox to chill for at least 10 minutes (alternatively 4C/wet ice can be used)

11.2. Pull enough Freezing Media A and Freezing Media B to create 1mL aliquots. The total mLs amount of freezing media needed is equal to the total number of aliquots needed.

12. Aspirate the supernatant

13. Resuspend the cells in Freezing Media A equal to one half of the total freezing media needed

14. Using a dropwise technique (1 drop/second) while swirling the sample, add Freezing Media B equal to the remaining half of the total volume.

15. Aliquot 1mL of cell suspension into each cryovial.

16. Place the cryovials into a CoolCell and into a -80° freezer for 24 hours (alternatively a Mr. Frosty or controlled rate freezer can be used)

17. Following this, immediately put the PBMCs cryovials into liquid nitrogen for long term storage

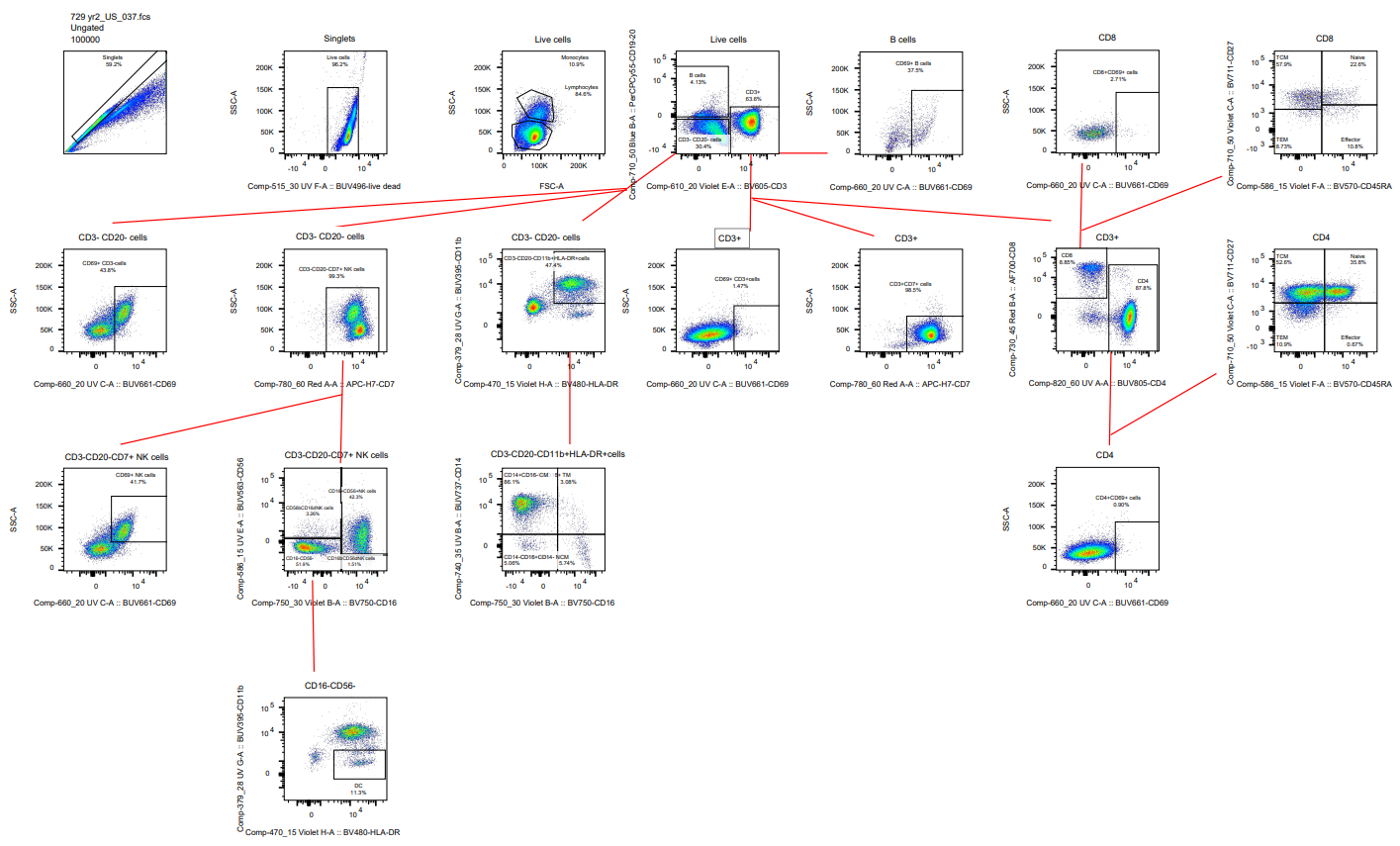

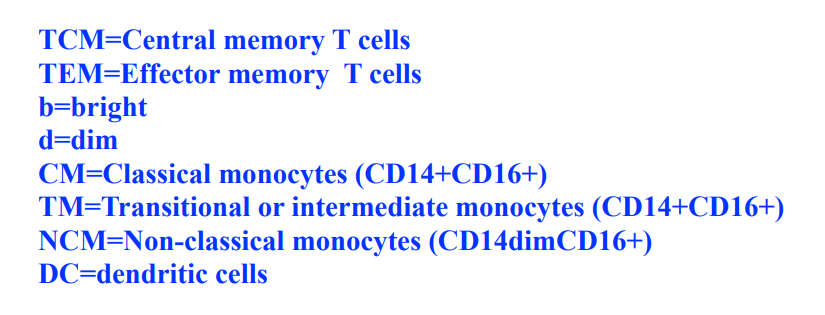

**Fig. S1.** Gating scheme for each cell type.

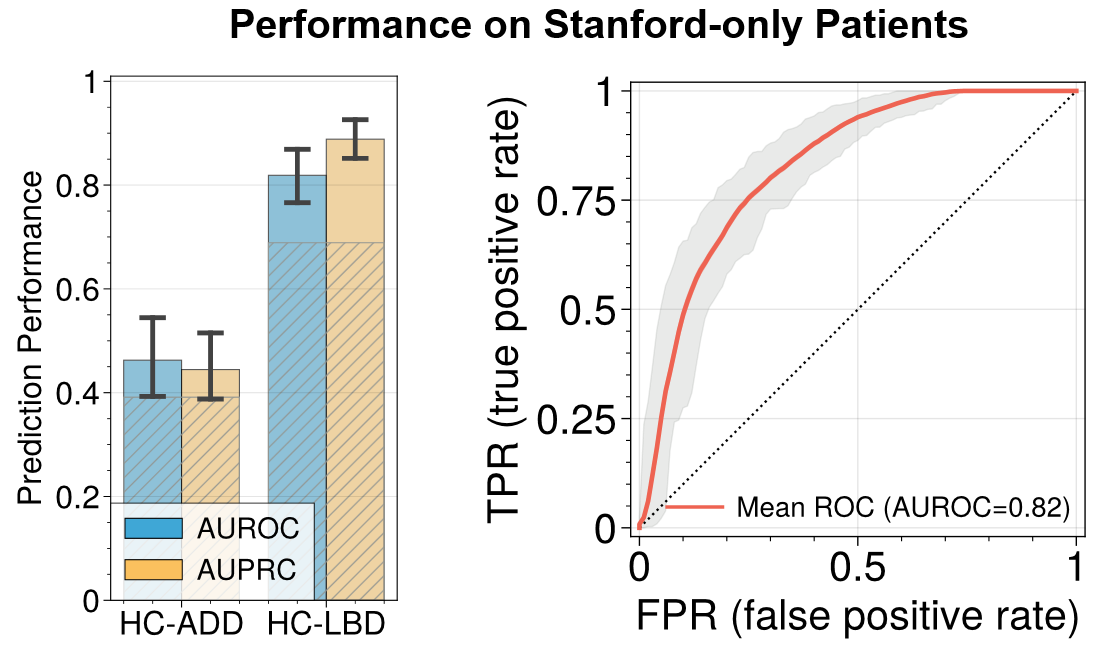

**Fig. S2.** Model performance based only on Stanford cohort retains high AUROC for HC-LBD with the ROC plotted on the right.

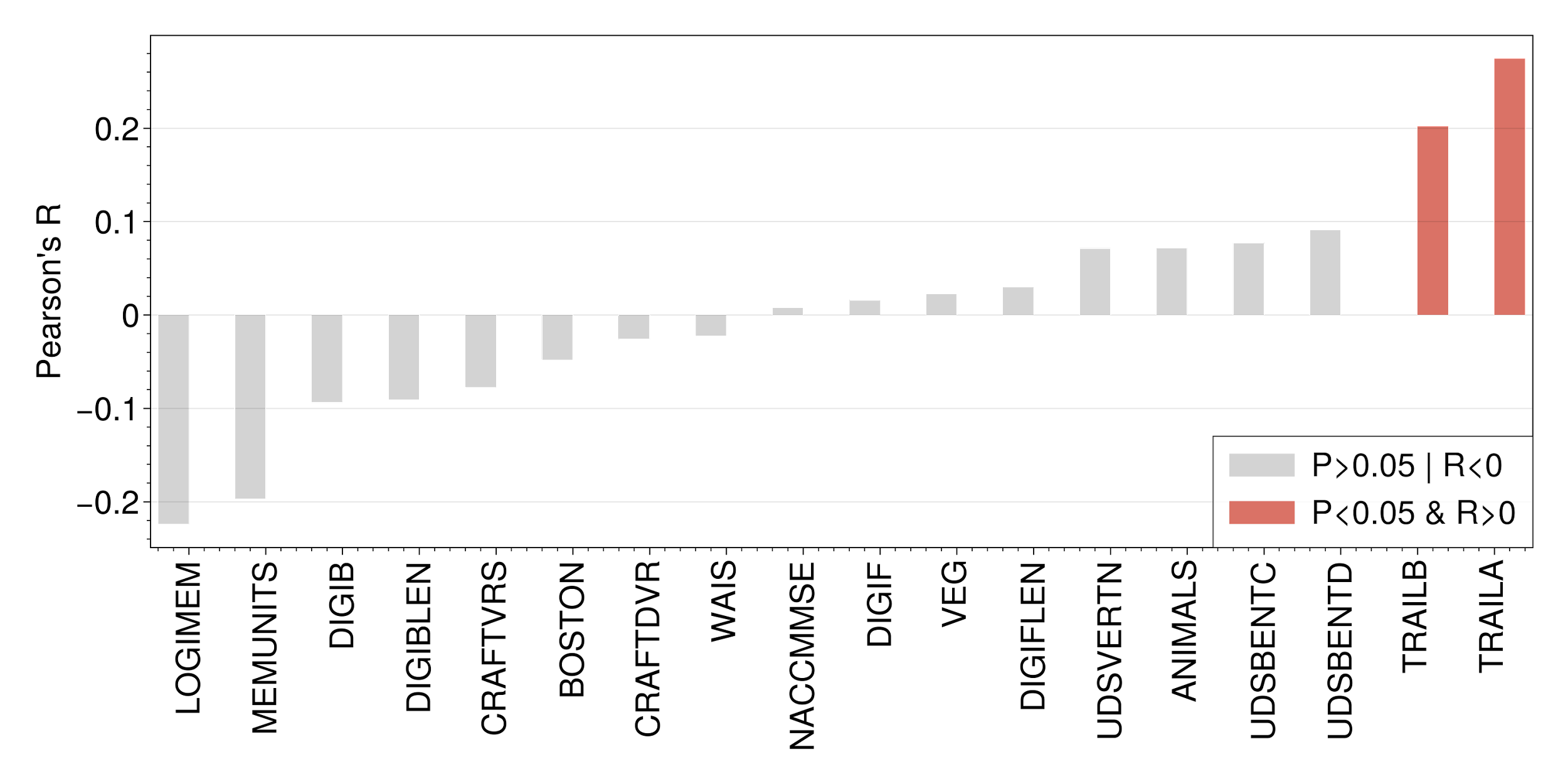

**Fig. S3.** Model performance (Pearson’s R) for the prediction of neuropsychological test scores in different cognitive domains. The acronym was taken directly from the NACC Data Dictionary available at: https://files.alz.washington.edu/documentation/uds3-rdd.pdf.

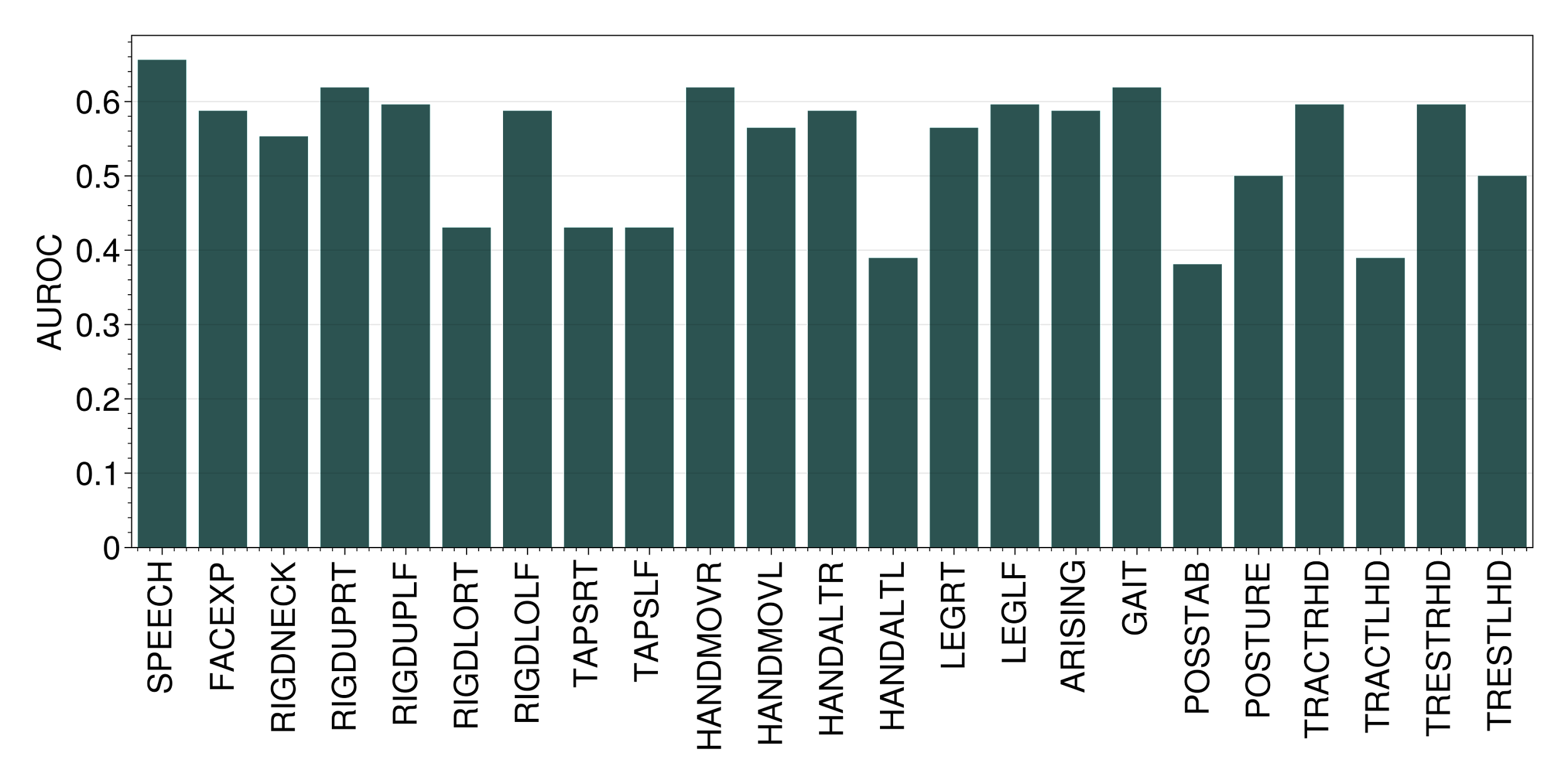

**Fig. S4.** Model performance (AUROC) for the prediction of motor examination. The acronym was taken directly from the NACC Data Dictionary available at: https://files.alz.washington.edu/documentation/uds3-rdd.pdf.

**Table S1.** Reagents used for flow cytometry experiments.

| **Stain Panel** | | | | |
| --- | --- | --- | --- | --- |
| **Filter** | **Fluorochrome** | **Marker** | **Catalog #** | **Clone** |
| 515/20 Blue D | AF488/FITC | pSTAT1 | BD 612596 | 4a |
| 710/50 Blue B | PerCPCy5.5 | CD20 | BD 558021 | H1 |
| 586/15 YG E | PE | p38 (pT180/pY182) | BD 612565 | 36/p38 |
| 610/20 YG D | PE-CF594 | Rab5 | sc-46692 AF594 | D-11 |
| 780/60 YG A | PE-Cy7 | pSTAT5 | BD 560117 | 47/Stat5(pY69) |
| 660/20 Red C | APC | pPLCγ2 | BD 558498 | K86-689.37 |
| 730/45 Red B | AF700/R718 | CD8 | BD 567354 | SK1 |
| 780/60 Red A | AF780 | CD7 | BD 564020 | M-T701 |
| 450/50 Violet I | BV421 | pS6 (V450) | BD 561457 | N7-548 |
| 470/15 Violet H | BV480 | HLA-DR | BD 566113 | G46-6 |
| 586/15 Violet F | BV570 | CD45RA | Biolegend 304132 | HI100 |
| 610/20 Violet E | BV605 | CD3 | Biolegend 300460 | UCHT1 |
| 710/50 Violet C | BV711 | CD27 | BD 564893 | M-T271 |
| 750/30 Violet B | BV750 | CD16 | BD 747145 | B73.1 |
| 800/30 Violet A | BV786 | CD107b (LAMP-2) | BD 565304 | H4B4 |
| 379/28 UV G | BUV395 | CD11b | BD 563839 | ICRF44 |
| 515/30 UV F | BUV496 | Live-Dead | L23105 |  |
| 586/15 UV E | BUV563 | CD56 | BD 612928 | NCAM16.2 |
| 660/20 UV C | BUV661 | CD69 | BD 750213 | FN50 |
| 740/35 UV B | BUV737 | CD14 | BD 612763 | M5E2 |
| 820/60 UV A | BUV805 | CD4 | BD 612887 | SK3 |
| **Stimulating Reagents** | | | | |
| **Stimulation** | **Catalog #** | **Company** | | |
| LPS | L7770 | Sigma -Aldrich | | |
| IFN𝜶 (alpha 2b) | 11105-1 | pbl assay science | | |
| IL-6 | BD 550071 | BD BIOSCIENCES | | |
| **Chemical Reagents** | | | | |
| **Chemical** | **Catalog #** | **Company** | | |
| Benzonase | 9025-65-4 | Signa | | |
| 1xPBS | SH30028.02 | Hyclone | | |
| FBS | SH30028.01 | Hyclone | | |
| RPMI-1640 | SH30027.01 | Hyclone | | |
| PFA | 4368 | Alfa Aesar | | |
| Methanol | A452SK-1 | Thermo Fisher Scientific | | |
